# Temporo-parietal brain regions are involved in higher order object perception

**DOI:** 10.1101/2020.04.13.039495

**Authors:** Sophia Nestmann, Daniel Wiesen, Hans-Otto Karnath, Johannes Rennig

**Author notes:** Address for correspondence:* Hans-Otto Karnath, Center of Neurology, University of Tübingen, D-72076 Tübingen, Germany.

## Abstract

Lesions to posterior temporo-parietal brain regions are associated with deficits in perception of global, hierarchical shapes, but also impairments in the processing of objects presented under demanding viewing conditions. Evidence from neuroimaging studies and lesion patterns observed in patients with simultanagnosia and agnosia for object orientation suggest similar brain regions to be involved in perception of global shapes and processing of objects in atypical (‘non-canonical’) orientation. In a localizer experiment, we identified individual temporoparietal brain areas involved in global shape perception and found significantly higher BOLD signals during the processing of non-canonical compared to canonical objects. In a multivariate approach, we demonstrated that posterior temporo-parietal brain areas show distinct voxel patterns for non-canonical and canonical objects and that voxel patterns of global shapes are more similar to those of objects in non-canonical compared to canonical viewing conditions. These results suggest that temporo-parietal brain areas are not only involved in global shape perception but might serve a more general mechanism of complex object perception. Our results challenge a strict attribution of object processing to the ventral visual stream by suggesting specific dorsal contributions in more demanding viewing conditions.

**Highlights:**

- Posterior temporo-parietal brain areas in the TPJ region that are involved in global shape perception are significantly involved in object perception
- Individual global shape TPJ ROIs identified with a specific localizer experiment prefer objects in non-canonical over objects in canonical orientations
- Univariate activations and multivariate voxel patterns in global shape TPJ ROIs distinguish canonical and non-canonical object presentations

## Introduction

Functional neuroimaging studies in neurological patients and healthy subjects identified posterior temporo-parietal brain regions as neural correlates of visual integration of local features into a global shape (Himmelbach, Erb, Klockgether, Moskau, & Karnath, 2009; Huberle & Karnath, 2012; Rennig, Bilalić, Huberle, Karnath, & Himmelbach, 2013; Rennig, Himmelbach, Huberle, & Karnath, 2015; Weissman & Woldorff, 2005; Zaretskaya, Anstis, & Bartels, 2013). These findings are in good agreement with lesion patterns of patients suffering from simultanagnosia (Balslev et al., 2014; Friedman-Hill et al., 1995; Himmelbach et al., 2009; Huberle & Karnath, 2006; Luria, 1959), who exhibit deficits in the perception of hierarchical, global objects (Bálint, 1909; Wolpert, 1924). However, several studies also showed that these patients have remarkable deficits perceiving coherent objects presented under demanding and atypical viewing conditions (Cooper & Humphreys, 2000; Rennig & Karnath, 2016).

While neuronal correlates of object processing are commonly attributed to the ventral visual stream (Goodale et al., 1991; Ungerleider & Mishkin, 1982) it was demonstrated that objects presented in atypical or non-canonical viewing conditions elicited neuronal responses outside ventral brain areas (Jacobs et al., 2015; James et al., 2002; Kosslyn et al., 1994; Terhune et al., 2005). It was also demonstrated that lesions to inferior parietal and temporo-parietal brain areas were associated with impairments in recognition of object orientation (Best, 1917; Jacobs et al., 2015; Karnath, Ferber, & Bülthoff, 2000; Martinaud et al., 2016; Turnbull, Laws, & McCarthy, 1995; Turnbull, Beschin, & Della Sala, 1997). These lesion patterns widely overlap with areas associated with global shape perception (Himmelbach et al., 2009; Huberle & Karnath, 2012; Rennig, Bilalić, Huberle, Karnath, & Himmelbach, 2013) and simultanagnosia (e.g. Friedman-Hill et al., 1995; Luria, 1959).

These findings indicate a similar neuronal mechanism for the integration of local features into a global shape and object perception under atypical or demanding viewing conditions. In the present study, we tested if the temporo-parietal junction (TPJ) that has been identified as a neuronal correlate of global shape perception (e.g. Huberle & Karnath, 2012; Rennig et al., 2013, 2015) also is involved in processes of higher order object perception. In a localizer fMRI experiment (Huberle & Karnath, 2012), we identified voxels in the anatomically defined TPJ area that were active during perception of global, hierarchical objects. In our main fMRI experiment, we tested if these TPJ voxels showed stronger signals during perception of objects presented in demanding or atypical (‘non-canonical’) orientations compared to objects presented in typical (‘canonical’) orientations. However, previous studies used different definitions of ‘canonical’ and ‘non-canonical’ viewing conditions varying from ‘frontal’ views (using regularly oriented stimuli for canonical views and images tilted for 90° or 180° for a non-canonical view; *e.g.* Best, 1917; Friedman-Hill et al., 1995; Karnath et al., 2000; Solms, Kaplan-Solms, Saling, & Miller, 1988; Turnbull et al., 1997, 1995) to ‘in-depth rotations’ (using the classic in-depth rotated ‘canonical’ view point and atypical in-depth rotations as ‘non-canonical’; *e.g.* Landau, Hoffman, & Kurz, 2006; Schendan & Stern, 2008; Terhune et al., 2005). A recent patient study also suggested different neuronal representations for frontal and in-depth views (Martinaud et al., 2016). To take this evidence into account, we created four different stimulus categories for our main fMRI experiment: objects in ‘frontal’ and ‘in-depth rotated’ views both as ‘canonical’ and ‘non-canonical’ conditions. This allowed us to explore if the TPJ shows specific responses to different kinds of viewing conditions.

## Methods

Twenty healthy volunteers *(mean* age□ = □25.65 years; *SD*□ =□3.91 years, 10 female, 1 lefthanded) participated in the study after giving written informed consent. They reported no history of neurological or psychiatric disorders. The experiment was approved by the ethics committee of the medical faculty of the University in Tübingen and conducted in accordance with the declaration of Helsinki. All participants had normal or corrected to normal vision.

Each participant was scanned using a 3T Siemens Magnetom TrioTim MRI system (Siemens AG, Erlangen, Germany) equipped with a 64-channel head coil. During a scanning session, participants performed two independent fMRI experiments. Stimuli were presented using Matlab (The Mathworks, Inc., Natick, MA, USA) and the Psychophysics Toolbox (Brainard, 1997; Pelli, 1997) and projected onto an MR compatible screen placed behind the bore of the scanner which could be viewed by the participants via a mirror mounted on the head coil. Behavioral responses were collected using a fiber-optic button response pad (Current Designs, Haverford, PA, USA) and eye movements were recorded during scanning using the Eye Link 1000 (SR Research Ltd., Ottawa, Ontario, Canada) with a sampling rate of 500 Hz.

In the first fMRI experiment, participants were presented with computer-generated object stimuli that were taken from the Object Databank (http://wiki.cnbc.cmu.edu/Objects) and from 3dcadbrowser (https://www.3dcadbrowser.com). Since there are several definitions of canonical and non-canonical object views (e.g. Blanz, Tarr, & Bülthoff, 1999; Verfaillie & Boutsen, 1995) we presented objects in four different viewing conditions (Figure 1A): *canonical rotated indepth view* (‘classic canonical view’, Blanz et al., 1999; Bülthoff, Edelman, & Tarr, 1995), *canonical frontal view, non-canonical in-depth view* and *non-canonical frontal view.* The experiment was carried out in four blocks, each consisting of 144 stimuli (36 of each viewing condition) presented in pseudo-random order (duration of each block: about 5.5 minutes). The stimulus selection was optimized to include as few similar objects as possible (*e.g.* to avoid too many car images in the stimulus sample) and to control for semantic frequency differences between *canonical* and *non-canonical* conditions. Semantic frequencies of German words were taken from the Leipzig Corpora Collection (Goldhahn et al., 2012). We created 1000 random stimulus sets and selected the best four sets (one for each presentation block) balancing semantic frequencies and reducing the number of repetitions of similar objects. All stimulus images were transformed to grayscale; size and luminance of the images was adjusted throughout the stimulus sample. The objects were presented at a size of 5° visual angles. To ensure that participants were paying attention to the object stimuli we conducted a task independent from the objective of our study. We choose objects, which could be classified as ‘made from metal’ or ‘not made from metal’ and participants were instructed to press one of two buttons for metal or non-metal objects. Semantic frequencies of these categories were also balanced throughout the four stimulus sets.

**Figure 1.**
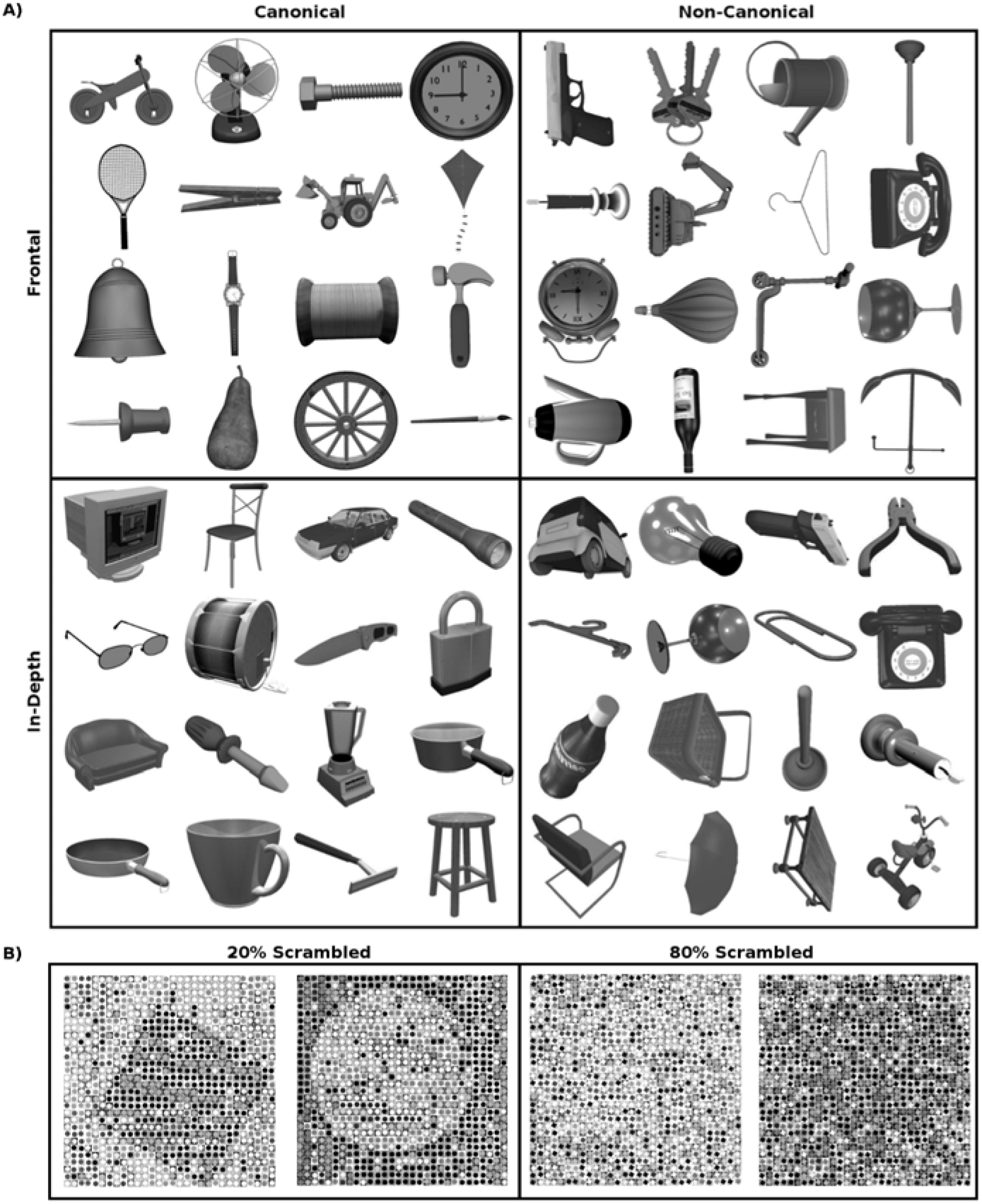
Stimulus material. **A)** In the first fMRI experiment, we presented objects from four different categories: objects in ‘frontal’ and ‘in-depth rotated’ views both as ‘canonical’ and ‘non-canonical’ conditions. **B)** In the second fMRI experiment, we showed the global shapes of either a circle or a square constructed from local images of squares or circles. All possible combinations of global and local elements were presented. The images were scrambled at two levels (20% and 80%) so that the global form could either be recognized (intact global perception) or not (scrambled global perception).

In the second fMRI experiment, we presented Navon-like global shape stimuli (Navon, 1977) that were applied in previous neuroimaging studies (Bloechle et al., 2018; Huberle & Karnath, 2012; Rennig et al., 2013, 2015). These stimuli showed the global shape of either a circle or a square constructed from local images of squares or circles and were presented in all possible combinations of global and local elements (congruent and incongruent). The stimulus images were scrambled at two levels (20% and 80%) so that the global form could either be recognized (‘intact global perception’) or not (‘scrambled global perception’, see Figure 1B). Stimuli were presented in two blocks (duration of each block: about 7 minutes), each consisting of 168 experimental trials (42 20%-circles, 42 20%-squares, 42 80%-circles, 42 80%-squares). The global forms were presented at 5.4° visual angles. Participants were instructed to respond via button press whether the stimulus was a circle or a square.

Both fMRI experiments were event-related designs and stimuli were presented for 300 ms with an inter-stimulus interval of 1700 ms. The events were ordered in an optimal rapid event-related design specified by optseq2 (Dale, 1999; https://surfer.nmr.mgh.harvard.edu/optseq). During the task, each participants’ eye position was tracked to control for alertness and stimulus fixation.

### MRI data acquisition

Functional images were acquired using multiband echo-planar-imaging (EPI) sequences. The first three participants were scanned using the following scanning parameters: TR□ = □2000 ms, TE□=□33 ms, flip angle □=□ 58°, FOV□=□1728 x 1728 mm^2^, 69 slices, voxel size□=□2□×□2□×□2 mm^3^, multiband factor = 3. All remaining participants were scanned with parameters for multiband EPI sequences from the HCP (Moeller et al., 2010): TR□=□1000 ms, TE□=□37 ms, flip angle □=□ 52°, FOV□=□1872 x 1872 mm^2^, 72 slices, voxel size□=□2□×□2□×□2mm^3^. Single band reference images (TR□ = □1000 ms, TE□=□37 ms, flip angle 52°, FOV□ = □1872 x 1872 mm^2^, 72 slices, voxel size□=□2□×□2□×□2 mm^3^) were collected before each functional run. T1-weighted anatomical scans (TR□=□2280 s, 176 slices, voxel size□ = □1.0□×□1.0□×□1.0 mm^3^; FOV□=□256 x 256 mm2, TE□ = □3.03 ms; flip angle□=□8°) were collected at the end of the experimental session.

### fMRI data analysis

Data pre-processing and model estimation was performed using SPM12 (http://www.fil.ion.ucl.ac.uk/spm). Functional images were realigned to each participants’ first image, aligned to the AC-PC axis and slice-time corrected. The original single-band image was then co-registered to the pre-processed functional images and the anatomical image was coregistered to the single-band image. The resolution of the single-band image was up-sampled before the anatomical image was aligned to it. Functional images were smoothed with a 4 mm FWHM Gaussian kernel. Time series of hemodynamic activation were modelled based on the canonical hemodynamic response function (HRF) as implemented in SPM12. Low-frequency noise was eliminated with a high-pass filter of 128 Hz. Correction for temporal autocorrelation was performed using an autoregressive AR(1) process. Movement parameters (roll, pitch, yaw; linear movement into x-, y-, z-directions) estimated in the realignment were included as regressors of no interest. To avoid inaccuracies associated with spatial normalization, analyses were conducted in subject space. For the first fMRI experiment, the experimental regressors consisted of the four experimental conditions: *canonical rotated in-depth view, canonical frontal view, non-canonical in-depth view* and *non-canonical frontal view.* A whole brain analysis of this experiment is presented in the supplementary material. For the second fMRI experiment, we used two experimental regressors: *intact global shapes* (20% scrambled) and *scrambled global shapes* (80% scrambled).

### Region of interest (ROI) analysis

Anatomical ROIs were created applying Freesurfer’s cortical reconstruction routine (Dale et al., 1999; Fischl et al., 2002) and the Destrieux atlas (Destrieux et al., 2010) for each subject. To create an individual anatomical TPJ ROI for each participant, we combined the posterior third of the superior temporal gyrus (Freesurfer Label 11174 and 12174), the sulcus intermedius primus (Freesurfer Label 11165, 12165), the angular gyrus (Freesurfer Labels 11125, 12125) and the posterior half of the supramarginal gyrus (Freesurfer Label 11126, 12126).

In a next step, we identified individual voxels that showed higher signals for 20%-scrambled global shapes compared to baseline as functional ROIs involved in global shape perception. The voxel-level threshold was set to *p* < 0.05 (uncorr.) without cluster threshold. Each participant’s individual global shape TPJ ROI was created as an intersection between the functional 20% scrambled *vs.* baseline contrast and the anatomical TPJ ROI. We were able to identify functional global shape TPJ ROIs in all of our 20 participants in the left and right hemisphere. Example structural and functional TPJ ROIs are presented in Figure 2A. The average size of the individual TPJ global shape ROIs was 2256.8 mm^3^ (SD = 2102 mm^3^) in the left hemisphere and 3049.2 mm^3^ (SD = 2569.5 mm^3^) in the right hemisphere. The mean center of mass was located at the MNI coordinates x = −43.41 (SD = 4.97); y = −56.36 (SD = 10.05); z = 32.52 (SD = 9.80) for left hemispheric global shape TPJ ROIs and x = 45.73 (SD = 6.28); y = – 52.66 (SD = 7.24); z = 35.49 (SD = 8.47) for right hemispheric global shape TPJ ROIs.

**Figure 2.**
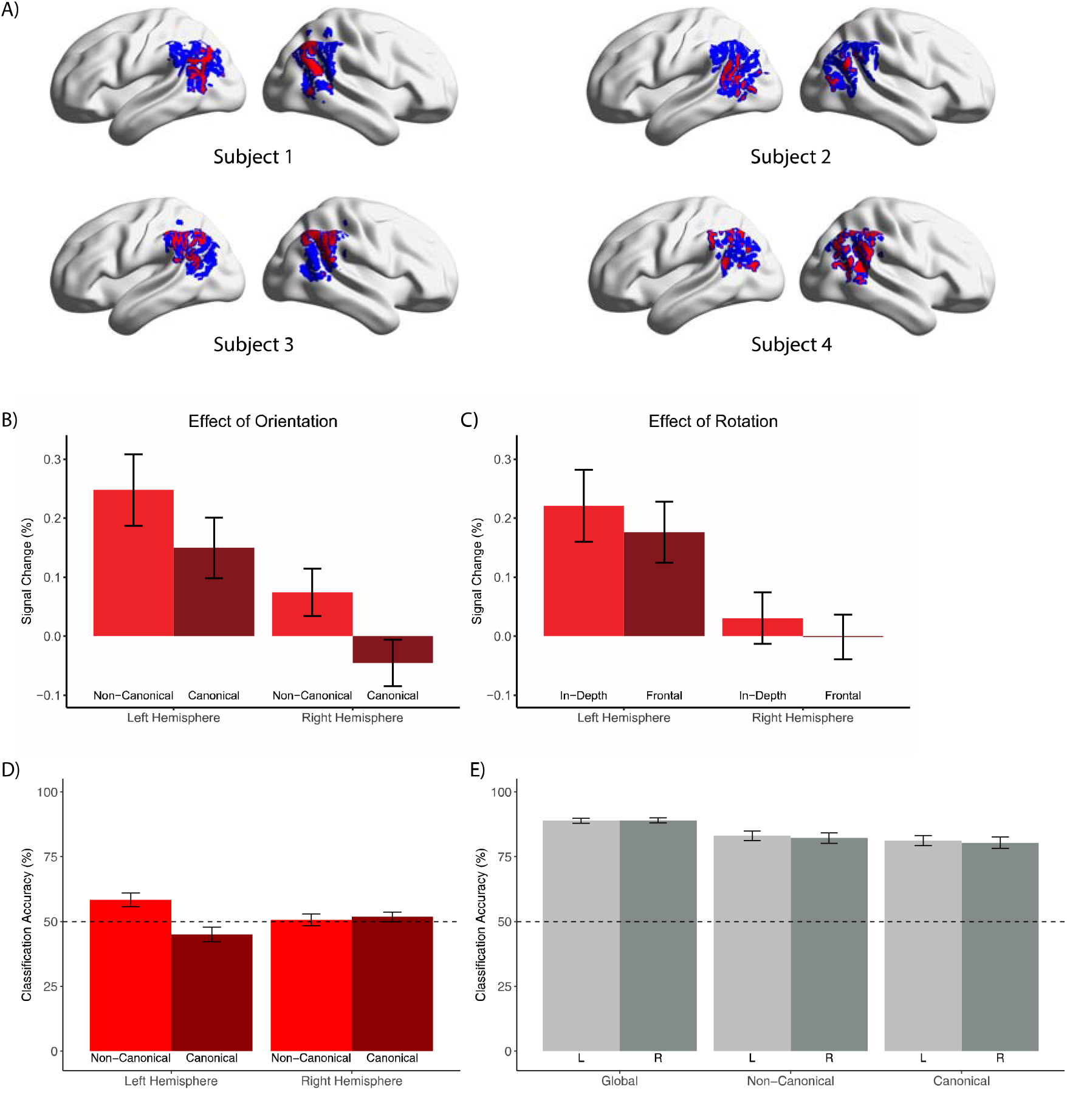
Univariate ROI Analysis and Multivoxel Pattern Analysis (MVPA). To analyze our data we created individual ROIs from which we extracted the mean percent signal change. **A)** Individual anatomical TPJ ROIs were created applying Freesurfer’s cortical reconstruction routine (Dale et al., 1999; Fischl et al., 2002) and the Destrieux atlas (Destrieux et al., 2010). From these ROIs, we then identified voxels that showed higher signals for 20%-scrambled global shapes compared to baseline for each subject. Percent signal change values and beta-coefficients extracted from these individual ROIs were then used for univariate and multivariate statistical analyses. Example ROIs from four representative subjects are presented in in standard MNI space on a surface version of the ch2 brain using the BrainNet viewer (Xia et al., 2013). **B)** *Effect of Orientation:* Average percent signal change values for non-canonical and canonical object presentations from left and right hemispheric global shape TPJ ROIs. There was a significant main effect for *orientation χ^2^* = 17.31, 3.17 × 10^-5^) and no significant interaction. **C)** *Effect of Rotation*: Average percent signal change values for in-depth and frontal object presentations from left and right hemispheric global shape TPJ ROIs. There was no significant main effect for *rotation* (*χ^2^* = 2.18,*p* = 0.140) and no significant interaction. **D)** *Classification of non-canonical and canonical voxel patterns in global shape TPJ ROIs:* Average classification accuracy (calculated as the proportion of correctly classified trials relative to all tested trials for both object viewing conditions) for non-canonical and canonical objects in left and right hemispheric global shape TPJ ROIs. Only non-canonical objects in the left hemispheric global shape TPJ ROIs were classified correctly above chance (*t_(19)_* = 3.23, *p* = 0.004). A linear mixed-effects model showed a significant interaction of orientation (non-canonical, canonical) and hemisphere (left, right) (*χ*^2^ = 9.43, *p* = 0.002). Subsequent paired *t*-tests showed a significant difference between classification accuracies for non-canonical and canonical objects in the left hemisphere (*t_(19)_* = 2.55, *p* = 0.019) but not in right hemisphere (*t_(19)_* = −0.39, *p* = 0.703). The dashed line indicates the 50% chance level. **E)** *Neuronal similarity of global shapes with non-canonical and canonical objects:* Average classification accuracy for global shapes, non-canonical and canonical objects in left and right hemispheric global shape TPJ ROIs. We calculated classification accuracy as the proportion of trials from every condition that were classified as global shapes relative to all tested trials from the respective condition. Classification accuracies of all object types from both hemispheres were classified above chance level as global shapes. Paired *t*-tests combining data from both hemispheres showed a significant difference between classification accuracies for global shapes and non-canonical (*t_(39)_* = 6.14, *p* = 3.2 × 10^-7^) as well as canonical objects (*t_(39)_* = 7.62, *p* = 3.0 × 10^-9^). There was also a significant difference between classification accuracies non-canonical and canonical objects (*t_(39)_* = 5.17, *p* = 7.2 × 10^-6^). The dashed line indicates the 50% chance level.

We used MarsBar (http://marsbar.sourceforge.net) to extract the mean percent signal change from these individual global shape TPJ ROIs for all four experimental conditions of the first fMRI experiment. For statistical data analysis we applied linear mixed effect models using R’s *lme4* and *lmerTest* packages. Model estimation was done using Restricted Maximum Likelihood (REML) estimation. Statistical significance was assessed using the *Anova* function provided by the *car* package.

## Results

### Behavioral Data

To ensure that participants were paying attention to the object stimuli, during the first experiment, participants were instructed to indicate via button press, whether an object was rather made from metal or not. For the first eleven participants’ we experienced technical problems for the collection of button presses. For participants with technical problems we were able to record responses in 84% of trials, those participants without technical problems responded in 93% of trials. We calculated a mixed-effect model to assess whether the percentage of correct answers depended on object orientation or rotation. Fixed effects were *orientation* (canonical *vs*. non-canonical orientation) and *rotation* (in-depth rotation *vs*. frontal views). The two subgroups of participants (with and without technical problems) were accounted for by including the fixed effect *group. Participant* was set as a random effect. We observed no significant main effect for *group* (*χ^2^* = 1.46, *p* = 0.227) and *rotation* (*χ^2^* = 2.76, *p* = 0.097). For the factor *orientation* we found a significant main effect (*χ^2^* = 5.14, *p* = 0.023) indicating that participants classifications were more often correct for canonical objects (84%) than for non-canonically presented views (81%). We also observed a significant interaction of the factors *orientation* and *group* (*χ^2^* = 5.53, *p* = 0.019; group *without* technical problems: canonical: 86% correct, non-canonical: 82%; group *with* technical problems: canonical: 81%, non-canonical: 81%). Canonically presented objects were classified correctly more often in the subgroup of participants without technical problems. None of the remaining interactions were significant (*p* ≥ 0.119). During the second experiment, participants were instructed to indicate, whether a stimulus showed a circle or a square. For participants with technical problems we were able to record responses in 82% of trials, those participants without technical problems responded in 89% of trials. For the presentation of 20% scrambled objects, answers were correct in 96%, for 80% scrambled objects in 52%.

### Univariate ROI Analysis

We observed stronger BOLD signals for non-canonical objects (see Figure 2B; mean percent signal change left hemisphere: 0.25%; mean right hemisphere: 0.07%) compared to canonical objects (mean left hemisphere: 0.15%; mean right hemisphere: −0.05%) in global shape TPJ ROIs. Splitting our data set by rotation we found slightly higher signals for in-depth rotated objects (see Figure 2C; mean left hemisphere: 0.22%; mean right hemisphere: 0.03%) compared to frontal objects (mean left hemisphere: 0.18%; mean right hemisphere: 0.00%). To statistically compare mean percent signal change values between conditions we calculated a linear mixed-effects model with fixed effects for *orientation* (canonical *vs.* non-canonical orientation), *rotation* (in-depth rotation *vs.* frontal views) and *hemisphere* (left *vs*. right) and *participant* as a random effect (Table 1). We observed a significant main effect for *orientation* (*χ^2^* = 17.31, *p* = 3.2 × 10^-5^) but not for *rotation* (in-depth rotated: 0.13%, frontal views: 0.09%; *χ^2^* = 2.18, *p* = 0.140) but we observed higher activations in the left (mean: 0.20%), compared to the right hemisphere (mean: 0.01%; *χ^2^* = 49.65, *p* = 1.8 × 10^-12^). No interactions were significant (for the full model output see Table 1).

**Table 1.** Parameter estimates and results of the univariate ROI analysis

|  | Estimate | Std.<br>Error | *df* | *t* | *p* |
| --- | --- | --- | --- | --- | --- |
| Fixed effects |  |  |  |  |  |
| Intercept | 0.108 | 0.070 | 33.00 | 1.55 | 0.130 |
| Hemisphere | 0.167 | 0.052 | 133.00 | 3.20 | 0.002 |
| Rotation | -0.068 | 0.052 | 133.00 | -1.29 | 0.199 |
| Orientation | -0.155 | 0.052 | 133.00 | -2.96 | 0.004 |
| Hemisphere x Rotation | 0.012 | 0.074 | 133.00 | 0.16 | 0.870 |
| Hemisphere x Orientation | 0.047 | 0.074 | 133.00 | 0.63 | 0.530 |
| Rotation x Orientation | 0.071 | 0.074 | 133.00 | 0.96 | 0.340 |
| Hemisphere x Rotation x Orientation | -0.050 | 0.105 | 133.00 | -0.48 | 0.631 |
| Random effect |  |  |  |  |  |
|  | Variance |  |  |  |  |
| Participant | 0.070 |  |  |  |  |
| Main effects |  |  |  |  |  |
| | $\chi^2$ | $p$ | | | |
| Hemisphere | 49.65 | $1.84 \times 10^{-12}$ | | | |
| Rotation | 2.18 | 0.140 |  |  |  |
| Orientation | 17.31 | $3.17 \times 10^{-5}$ | | | |
| Interactions |  |  |  |  |  |
| Hemisphere x Rotation | 0.06 | 0.804 |  |  |  |
| Hemisphere x Orientation | 0.17 | 0.682 |  |  |  |
| Rotation x Orientation | 0.77 | 0.312 |  |  |  |
| Hemisphere x Rotation x Orientation | 0.23 | 0.631 |  |  |  |

We extracted the mean percent signal change values of canonical and non-canonical objects in frontal and in-depth rotated views from individual global shape TPJ ROIs. *Orientation*, *hemisphere* and *rotation* were set as fixed effects and *participant* was set as random effect in a linear mixed-effects model. The upper rows show the parameter estimates for each factor in the model as well as the variance of the random effect. The lower rows show the results of the chi-square test of the model, with one degree of freedom for each factor.

Since we observed a significant effect between canonical and non-canonical object orientations in our behavioral data (see above) we were interested if the significant effect in BOLD responses between canonical and non-canonical object orientations can be explained by behavior *(i.e.* task difficulty). To statistically assess the influence of individual behavior on BOLD responses we calculated a linear mixed-effects model with fixed effects for *orientation* (canonical *vs.* non-canonical orientation), *percent correct* (percent of correctly classified objects as a continuous predictor) and *participant* as a random effect. We observed a significant main effect for *orientation* (*χ^2^* = 12.94, *p* = 3.2 × 10^-5^) but not for *percent correct* (*χ^2^*= 0.29, *p* = 0.591) and no significant interaction (*χ^2^* = 0.79, *p* = 0.375).

### Multivoxel Pattern Analysis (MVPA)

While a univariate analysis can demonstrate differences in signal strengths of experimental conditions in an ROI, *e.g.* stronger signals for non-canonical views *vs.* canonical views in TPJ regions, a MVPA is able to quantify neuronal representational similarities between experimental conditions (Haxby et al., 2001; Haynes & Rees, 2006). First, we created feature vectors from fMRI data by applying the approach suggested by Mumford et al. (2012): for both experiments (objects in canonical and non-canonical viewing conditions, global shapes) and every participant we calculated beta regression coefficient images for each experimental trial separately by running a general linear model including a regressor for the respective trial as well as another regressor for all other trials. For this analysis we used unsmoothed images and did not apply any high-pass filtering in the statistical model. Resulting beta values of voxels from individual global shape TPJ ROIs were then used as features for training and testing support vector machines (SVM). The *R* package *e1071* was used to train SVMs.

In a first analysis, we aimed at showing that TPJ areas responding to intact global shapes show specific voxel pattern responses for canonical and non-canonical object views. The prediction was that if TPJ areas responding to intact global shapes show stronger univariate activations for processing of non-canonical compared to canonical object views these TPJ areas should also show different voxel patterns for these two object viewing conditions. Per participant and hemisphere, we selected beta values for every experimental trial from the first fMRI experiment (canonical and non-canonical object views) from every voxel of the previously defined global shape TPJ ROI. Each experimental trial was treated as an observation and each voxel as a feature for the machine learning model. We split the data into a training set (80% of trials) and a test set (20% of trials) and trained a SVM with a radial basis kernel with the training set. We conducted grid search using 10-fold cross validation to optimize the regularization parameter C = [0.01, 0.1, 1, 10, 100] and gamma = [0.1, 0.5, 1, 2]. Using the SVM model, we predicted from the voxel patterns of the test set (per participant and hemisphere) if an individual trial was a canonical or a non-canonical object. We calculated classification accuracy as the proportion of correctly classified trials relative to all tested trials for both object viewing conditions. In the left hemisphere, we observed classification accuracies above chance for non-canonical objects (Figure 2D; mean classification accuracy: 58.4%; *t*-test against 50% chance level: *t_(19)_* = 3.23, *p* = 0.004) but not for canonical objects (mean classification accuracy: 45.0%; *t*-test against 50% chance level: *t_(19)_* = −1.76, *p* = 0.093). In the right hemisphere, we did not observe classification accuracies above chance for non-canonical objects (mean classification accuracy: 50.6%; *t*-test against 50% chance level: *t_(19)_* = 0.27, *p* = 0.790) and canonical objects (mean classification accuracy: 51.8%; *t*-test against 50% chance level: *t_(19)_* = −0.94, *p* = 0.357). To asses statistical differences of SVM classification between object viewing conditions and hemispheres we used individual classification accuracies as a dependent variable in a linear mixed-effects model with fixed effects for *orientation* (canonical *vs.* non-canonical orientation) and *hemisphere* (left *vs.* right) and *participant* as a random effect. We found a significant main effect of orientation (*χ*^2^ = 6.70, *p* = 0.010) but not for hemisphere (*χ^2^* = 0.05, *p* = 0.826) and a significant interaction (*χ* = 9.43, *p* = 0.002). Subsequent paired *t*-tests showed a significant difference between classification accuracies for non-canonical and canonical objects in the left hemisphere (58.4% *vs.* 45%; *t_(19)_* = 2.55, *p* = 0.019) but not in right hemisphere (50.6% *vs*. 51.8%; *t_(19)_* = −0.39, *p* = 0.703).

A second multivariate analysis aimed at showing similarity between processing of global shape stimuli and objects in non-canonical presentation conditions. The prediction is that a machine learning model trained on response patterns of global shape stimuli from the global shape TPJ should classify objects in non-canonical viewing conditions as more similar to global shapes than objects in canonical viewing conditions. First, we selected all intact global shape trials (20% scrambled) from the second fMRI experiment. Per participant and hemisphere, we selected 80% of these trials as training data. We trained a one-class SVM with a radial basis kernel (nu = 0.5) using a 10-fold cross validation for classification optimization with the global shape training set. To test classification accuracy we used the remaining 20% test global shape trials and all canonical and non-canonical object trials from the first fMRI experiment. We calculated classification accuracy as the proportion of trials from every condition (global shapes, canonical and non-canonical objects) that were classified as global shapes relative to all tested trials from the respective condition. In the left hemisphere, we observed classification accuracies above chance for global shapes (Figure 2E; mean classification accuracy: 88.8%; *t*-test against 50% chance level: *t_(19)_* = 41.65, *p* = 3.9 × 10^-20^), non-canonical objects (mean classification accuracy: 83.0%; *t*-test against 50% chance level: *t_(19)_* = 18.01, *p* = 2.1 × 10^-12^) and canonical objects (mean classification accuracy: 81.2%; *t*-test against 50% chance level: *t_(19)_* = 16.57, *p* = 9.4 × 10^-13^). In the right hemisphere, classification accuracies were above chance for global shapes (Figure 2D; mean classification accuracy: 89.0%; *t*-test against 50% chance level: *t_(19)_* = 41.33, *p* = 4.5 × 10^-20^), non-canonical objects (mean classification accuracy: 82.3%; *t*-test against 50% chance level: *t_(19)_* = 15.89, *p* = 2.0 × 10^-13^) and canonical objects (mean classification accuracy: 80.4%; *t*-test against 50% chance level: *t_(19)_* = 14.11, *p* = 1.6 × 10^-11^). To assess statistical differences of SVM classification between conditions and hemispheres we used individual classification accuracies as a dependent variable in a linear mixed-effects model with fixed effects for *object type* (global shapes *vs.* canonical *vs.* non-canonical orientation) and *hemisphere* (left *vs.* right) and *participant* as a random effect. We found a significant main effect of object type = 6.70, *p* (*χ^2^* = 4.1 × 10^-9^) but not for hemisphere (*χ^2^* = 0.19, *p* = 0.658) and no significant interaction (*χ^2^* = 0.15, *p* = 0.927). Subsequent paired *t*-tests combining data from both hemispheres showed a significant difference between classification accuracies for global shapes and non-canonical (88.9% vs. 82.6%; *t_(39)_* = 6.14, *p* = 3.2 × 10^-7^) as well as canonical objects (88.9% vs. 80.8%; *t_(39)_* = 7.62, *p* = 3.0 × 10^-9^). However, there was also a significant difference between classification accuracies of non-canonical and canonical objects (82.6% vs. 80.8%; *t_(39)_* = 5.17, *p* = 7.2 × 10^-6^) indicating that non-canonical objects were significantly more often classified as a global shape than canonical objects.

### Control analysis

As control ROI we took all voxels of the individual, anatomical TPJ ROIs not responding to global shapes. The average size of these individual control ROIs was 12766 mm^3^ (SD = 2204 mm^3^) in the left hemisphere and 11364 mm^3^ (SD = 3705 mm^3^) in the right hemisphere. The mean center of mass was located at the MNI coordinates x = −44.53 (SD = 3.14); y = −59.87 (SD = 5.72); z = 29.53 (SD = 6.85) for left hemispheric and x = 49.15 (SD = 3.99); y = −51.20 (SD = 6.67); z = 30.32 (SD = 6.01) for right hemispheric ROIs.

In contrast to our global shape TPJ ROIs we observed significant deactivations in the remainder of the anatomically defined TPJ. We found negative BOLD signals for non-canonical objects (mean percent signal change left hemisphere: −0.19%; mean right hemisphere: −0.15) and canonical objects (mean left hemisphere: −0.27%; mean right hemisphere: −0.24%). Accordingly, we observed deactivations for objects presented in in-depth rotations (mean left hemisphere: – 0.22%; mean right hemisphere: −0.19%) and frontal views (mean left hemisphere: −0.24%; mean right hemisphere: −0.20%). We calculated a linear mixed effects model with percent signal change values as dependent variable, fixed effects for *orientation* (canonical vs. non-canonical orientation)*, rotation* (in-depth rotation vs. frontal views) and *hemisphere* (left vs. right) and *participant* as a random effect (Table 2). We observed a significant main effect for *orientation* = 20.93, 4.76 × 10^-6^).

**Table 2.**
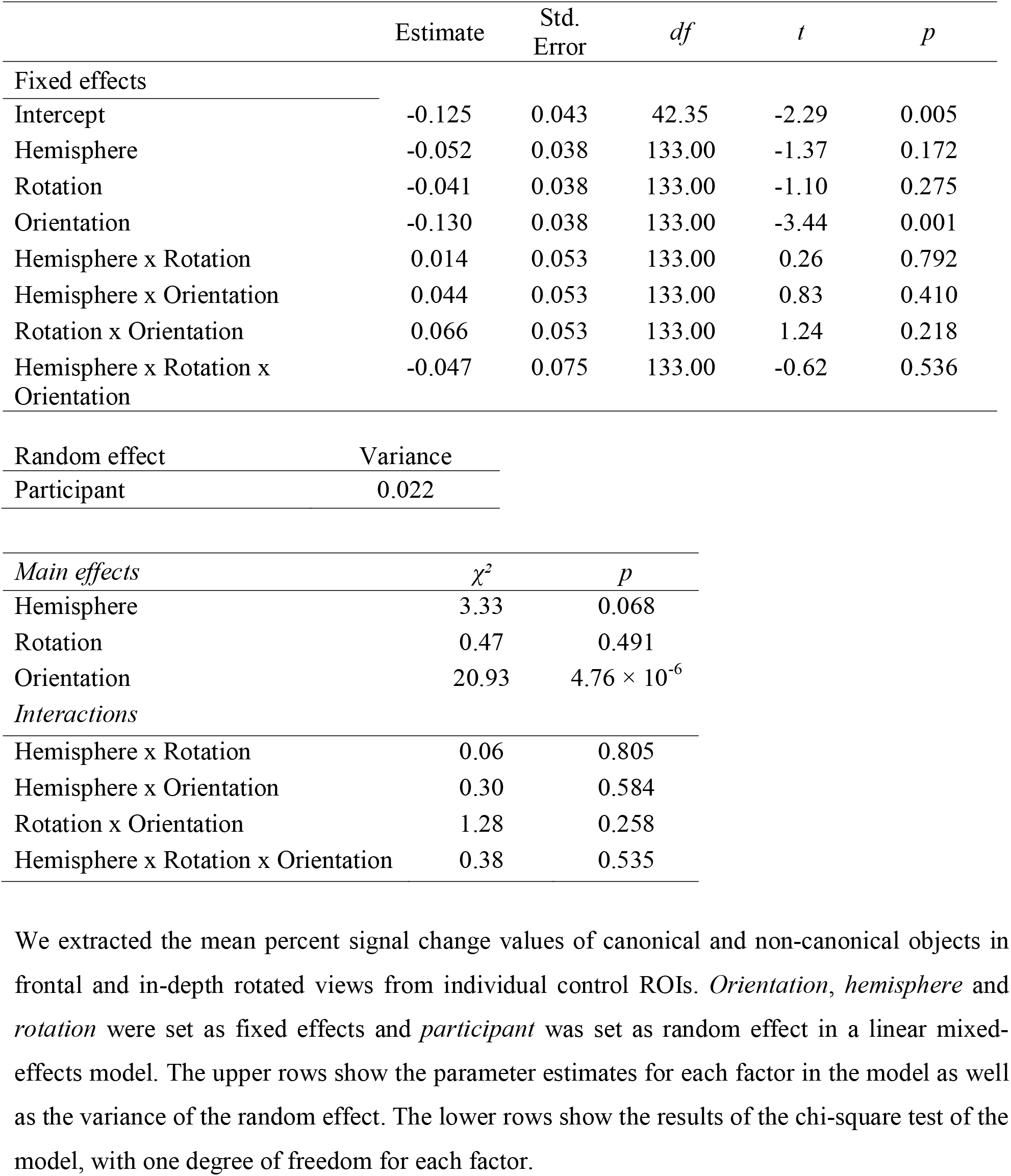
Parameter estimates and results of the control analysis

|  | Estimate | Std.<br>Error | *df* | *t* | *p* |
| --- | --- | --- | --- | --- | --- |
| **Fixed effects** |  |  |  |  |  |
| Intercept | -0.125 | 0.043 | 42.35 | -2.29 | 0.005 |
| Hemisphere | -0.052 | 0.038 | 133.00 | -1.37 | 0.172 |
| Rotation | -0.041 | 0.038 | 133.00 | -1.10 | 0.275 |
| Orientation | -0.130 | 0.038 | 133.00 | -3.44 | 0.001 |
| Hemisphere x Rotation | 0.014 | 0.053 | 133.00 | 0.26 | 0.792 |
| Hemisphere x Orientation | 0.044 | 0.053 | 133.00 | 0.83 | 0.410 |
| Rotation x Orientation | 0.066 | 0.053 | 133.00 | 1.24 | 0.218 |
| Hemisphere x Rotation x Orientation | -0.047 | 0.075 | 133.00 | -0.62 | 0.536 |
| **Random effect** |  |  |  |  |  |
|  | Variance |  |  |  |  |
| Participant | 0.022 |  |  |  |  |
| **Main effects** |  |  |  |  |  |
| | $\chi^2$ | *p* | | | |
| Hemisphere | 3.33 | 0.068 |  |  |  |
| Rotation | 0.47 | 0.491 |  |  |  |
| Orientation | 20.93 | $4.76 \times 10^{-6}$ | | | |
| **Interactions** |  |  |  |  |  |
| Hemisphere x Rotation | 0.06 | 0.805 |  |  |  |
| Hemisphere x Orientation | 0.30 | 0.584 |  |  |  |
| Rotation x Orientation | 1.28 | 0.258 |  |  |  |
| Hemisphere x Rotation x Orientation | 0.38 | 0.535 |  |  |  |

We extracted the mean percent signal change values of canonical and non-canonical objects in frontal and in-depth rotated views from individual control ROIs. *Orientation*, *hemisphere* and *rotation* were set as fixed effects and *participant* was set as random effect in a linear mixed-effects model. The upper rows show the parameter estimates for each factor in the model as well as the variance of the random effect. The lower rows show the results of the chi-square test of the model, with one degree of freedom for each factor.

## Discussion

The main research question of the present fMRI study was to investigate the role of the TPJ in processing of objects under demanding viewing conditions. First, we localized voxels in the anatomically defined TPJ area that were active during perception of intact global shapes (Huberle & Karnath, 2012). In our main experiment, we presented coherent objects in canonical and non-canonical views. We found that those TPJ areas that preferred global shapes showed higher BOLD signals during processing of objects in non-canonical compared to canonical presentation conditions. These results suggest that temporo-parietal brain areas might serve a more general mechanism of complex object perception. In a MVPA we were able to show that TPJ areas responding to global shapes have specific activation patterns for canonical and non-canonical objects. In a different machine learning approach, we could demonstrate that the activation pattern of global shapes is significantly more similar to non-canonical than canonical objects.

Our results offer an explanation for the observation that deficits in the perception of complex objects, like global shapes (Friedman-Hill et al., 1995; Himmelbach et al., 2009; Huberle & Karnath, 2006; Luria, 1959) and coherent objects in demanding viewing conditions (Cooper & Humphreys, 2000; Rennig & Karnath, 2016), as well as impairments for the perception of object orientation (Best, 1917; Jacobs et al., 2015; Karnath, Ferber, & Bülthoff, 2000; Martinaud et al., 2016; Turnbull, Laws, & McCarthy, 1995) occur after lesions to temporo-parietal brain areas. The present findings suggest a common mechanism for perception of objects that need additional visual processing steps compared to objects that are clearly visible and are primarily processed through the ventral visual stream (Goodale et al., 1991; Ungerleider & Mishkin, 1982). Our results are also in good agreement with functional neuroimaging studies (Sugio et al., 1999; Terhune et al., 2005) that found significant activations in temporo-parietal brain areas for perception of objects in non-canonical compared to canonical viewing conditions supporting the assumption that processing of non-canonical compared to canonical views might require additional visual processing steps. Our multivariate analyses support the assumption of a functional overlap of global shape perception and object perception in demanding viewing conditions in posterior temporo-parietal brain areas. We were able to demonstrate that global shape TPJ regions have specific activation patterns for non-canonical and canonical objects and that neuronal representation patterns of global shapes are significantly more similar to representations of non-canonical than canonical objects.

Neuronal signals along the dorsal visual stream were reported by several functional neuroimaging studies that investigated different questions on object perception (Dekker et al., 2011; Freud, Culham, et al., 2017; Konen & Kastner, 2008; Kourtzi & Kanwisher, 2000). In particular, it was demonstrated that areas along the dorsal visual stream are significantly involved in processing of man-made objects and tools (Chao & Martin, 2000; Kristensen et al., 2016; Mahon et al., 2013; Mruczek et al., 2013). Recent electrophysiological monkey and human neuroimaging studies also showed that perception of three-dimensional objects elicit activations particularly in dorsal regions (Freud et al., 2018; Freud, Ganel, et al., 2017; Janssen et al., 2018; Van Dromme et al., 2016). These findings support the results of the present study with significant BOLD signals in the TPJ during conditions of complex object presentations (canonical and non-canonical object stimuli) since especially objects in non-canonical viewing conditions may require processes of three-dimensional perception to enable successful object recognition.

In addition to more dorsally located brain areas in close vicinity to the TPJ, we observed inferior temporal brain regions along the ventral visual stream to show stronger BOLD signals for the perception of non-canonical compared to canonical objects. This is in line with results described by Terhune et al. (2005) and Sugio et al. (1999) that demonstrated significant clusters in dorsal and ventral areas for the contrast of non-canonical *vs.* canonical object stimuli. We observed clusters of activation in the LOC (Grill-Spector, Kushnir, Hendler, & Malach, 2000; Grill-Spector, Kourtzi, & Kanwisher, 2001) and the fusiform gyrus (Haxby et al., 2001).

Alivisatos and Petrides (1997) report left hemispheric posterior parietal brain areas to be involved in both the processing of mirrored stimuli and mental rotation. Our observation of increased activation in TPJ areas during perception of non-canonical views might also be interpreted as an effect of mental rotation. To address possible effects of mental rotation in our data, we tested if participants’ reaction times changed due to rotation angle of the object stimuli. If participants mentally rotated the objects before recognizing them, the response times should be higher with increasing degree of rotation (Shepard & Metzler, 1971). We analyzed reaction times to frontal images, which were presented at rotation angles of 90° and 180° from the canonical view (0°). We excluded 26 images of objects, which had an unambiguous upright orientation. In a one-way ANOVA we could not identify any significant differences in reaction times between the three orientations (*F* = 1.61, *p* = 0.209). This supports the interpretation that our results cannot be explained by mental rotation only. With short presentation times in a fast event-related design (300 ms), our study design was also significantly different from paradigms applied to study mental rotation using fMRI (Schendan & Stern, 2007, 2008) that used presentation times allowing active mental rotation of objects in non-canonical viewing conditions (1875 ms and 3750 ms). Our short presentation times with rapid, consecutive stimulus presentations hardly allowed for active mental rotation. However, contrasting the effect of mental rotation from perceptual processes in the TPJ should be subject of future research.

On a theoretical level, the present study combines two established concepts in cognitive psychology: Gestalt perception (Koffka, 1935; Wertheimer, 1923) and object constancy (Carlson, 1962). Gestalt perception describes mechanisms of higher object perception, like holistic integration of local elements into global entities (Navon, 1977), scene perception, visual grouping and object completion (Wagemans, Elder, et al., 2012; Wagemans, Feldman, et al., 2012). Object constancy on the other hand, describes mechanisms that allow us to recognize objects in atypical or demanding viewing conditions, for example from multiple angles or despite of occlusion or the presence of visual noise (Carlson, 1962). Our data indicates a significant overlap of these two otherwise independent concepts, especially considering that deficits in Gestalt perception (Friedman-Hill et al., 1995; Huberle & Karnath, 2006; Luria, 1959) and impairments in object constancy (Cooper & Humphreys, 2000; Rennig & Karnath, 2016) occur after comparable lesion patterns to temporo-parietal brain areas.

Our results challenge a strict attribution of object processing to the ventral visual stream by suggesting dorsal contributions in more demanding viewing conditions. In particular, we demonstrated contributions of TPJ areas in the perception of not only global shapes, but also objects presented in atypical viewing conditions. In line with previous findings (Cooper & Humphreys, 2000; Rennig & Karnath, 2016), our present observations suggest that temporoparietal brain areas serve a common mechanism for perception of objects that require additional visual processing steps to achieve higher order object representations.

## Acknowledgements

This work was supported by the Deutsche Forschungsgemeinschaft (RE 3693/1-1 to J.R.). We thank Lisa Röhrig for assistance during MRI scanning sessions.

## Supplementary Methods & Results

### Whole brain analysis

#### Methods

For the whole brain analysis functional and anatomical images were transferred to standard MNI space (Montreal Neurological Institute, McGill University, Montreal, Canada) using SPM. Functional images were smoothed with a 6 mm FWHM Gaussian kernel. The remaining preprocessing steps as well as model estimations were then performed as described for the analyses in native space (see *fMRI data analysis*). Within voxels significantly active throughout the whole experiment (Omnibus *F* contrast with *p* > 0.05 uncorr.) we calculated group-level contrasts to identify brain regions significantly involved in processing of objects presented in *canonical* and *non-canonical* or *frontal* or *rotated* orientations. The voxel-level threshold had to be set to *p* < 0.001 (uncorr.) with a cluster threshold of 100 voxels since no voxels survived a correction for multiple comparisons (FWE correction, *p* < 0.05). Identification of cortical areas from functional clusters was done using the MNI2TAL converter (https://bioimagesuiteweb.github.io/webapp/mni2tal.html).

#### Results

Clusters identified as showing higher activation for objects presented in non-canonical compared to canonical views (across in-depth and frontal views) were observed in the lateral occipital complex (LOC) of both hemispheres with the global maximum in the left hemisphere. Further clusters were found in the fusiform gyri of both hemispheres as well as in the right supplementary motor cortex, the right parietal cortex, the right inferior frontal cortex and the right dorsolateral prefrontal cortex (Table S1; Figure S1A). No clusters showing higher activation for canonical objects compared to non-canonically oriented objects survived.

Clusters identified as showing higher activation for objects presented in in-depth views compared to frontal views (across non-canonical and canonical views) were observed in the occipital cortices of both hemispheres with the global maximum in the left hemisphere. Additional clusters were found in the precentral cortices (including premotor and supplementary motor cortex), and frontal cortices of both hemispheres as well as in the left fusiform gyrus (Table S1; Figure S1B). No clusters survived the comparison of frontal views *vs.* in-depth rotations.

Next, we compared non-canonical *vs.* canonical objects in frontal views and identified significant clusters in the parietal lobes of both hemispheres, with the global maximum in the right superior parietal lobe. Additional clusters were found in the LOC, the premotor and supplementary motor cortex of both hemispheres, the right angular gyrus, the right fusiform, the right inferior frontal and the left frontal cortex (Table S1; Figure S1C). No clusters survived the reverse comparison of canonical orientation *vs.* non-canonical orientation.

For objects presented in in-depth rotated views, clusters identified as showing higher activation for objects presented in non-canonical orientations were located in the LOC of both hemispheres with the global maximum in the right hemisphere. Further clusters were found in the right superior parietal lobe (Table S1; Figure S1D). No clusters showing higher activation for canonical orientation compared to non-canonical orientation could be identified.

**Table S1.**
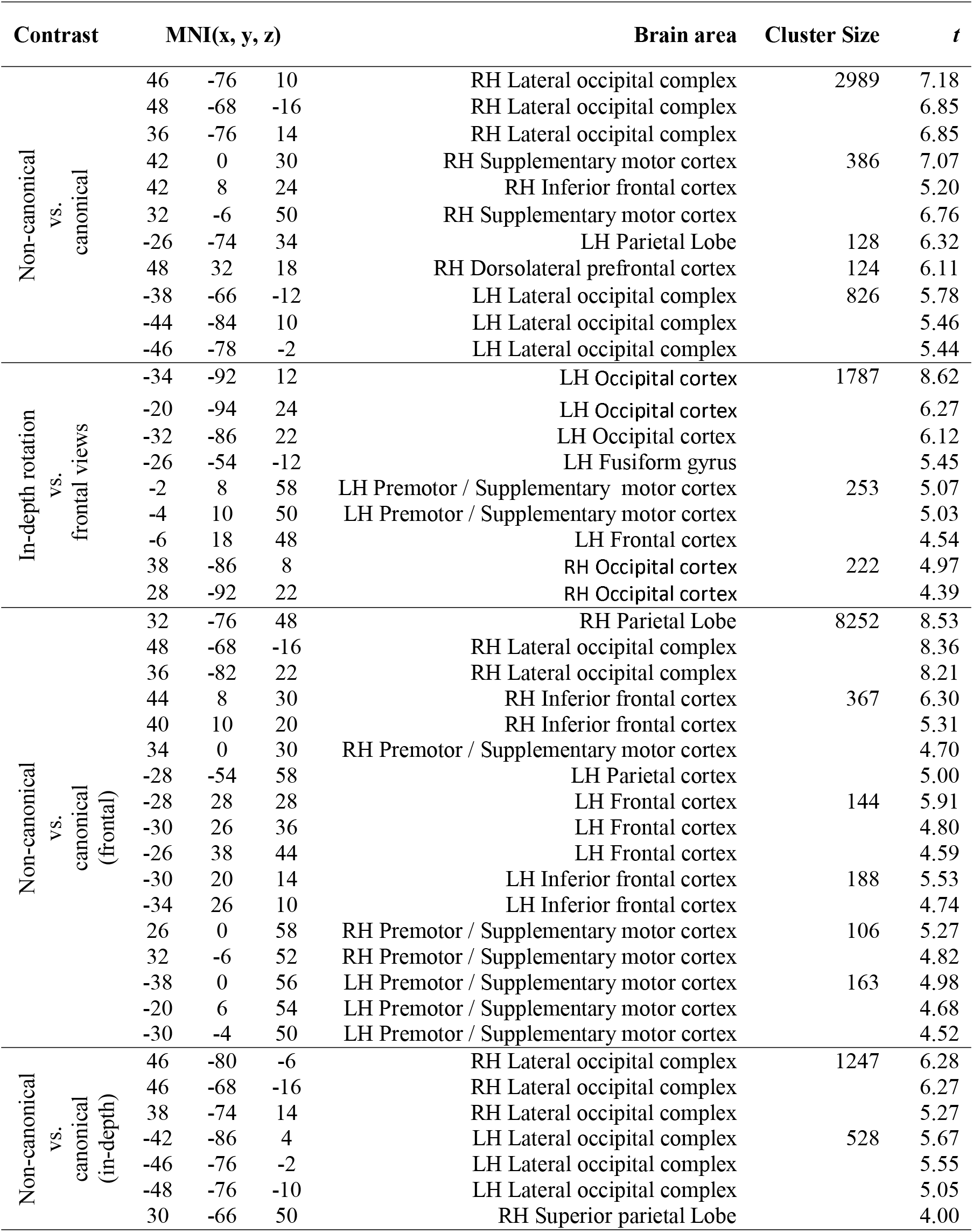
Brain areas assigned to the significant clusters of the contrasts non-canonical *vs.* canonical, in-depth rotation *vs.* frontal views, non-canonical *vs.* canonical for frontal views and non-canonical *vs.* canonical for in-depth rotated views. The table shows coordinates, cluster size and *t* values for all clusters of *p* ≤ 0.001 with a cluster threshold > 100 voxels.

**Figure S1.**
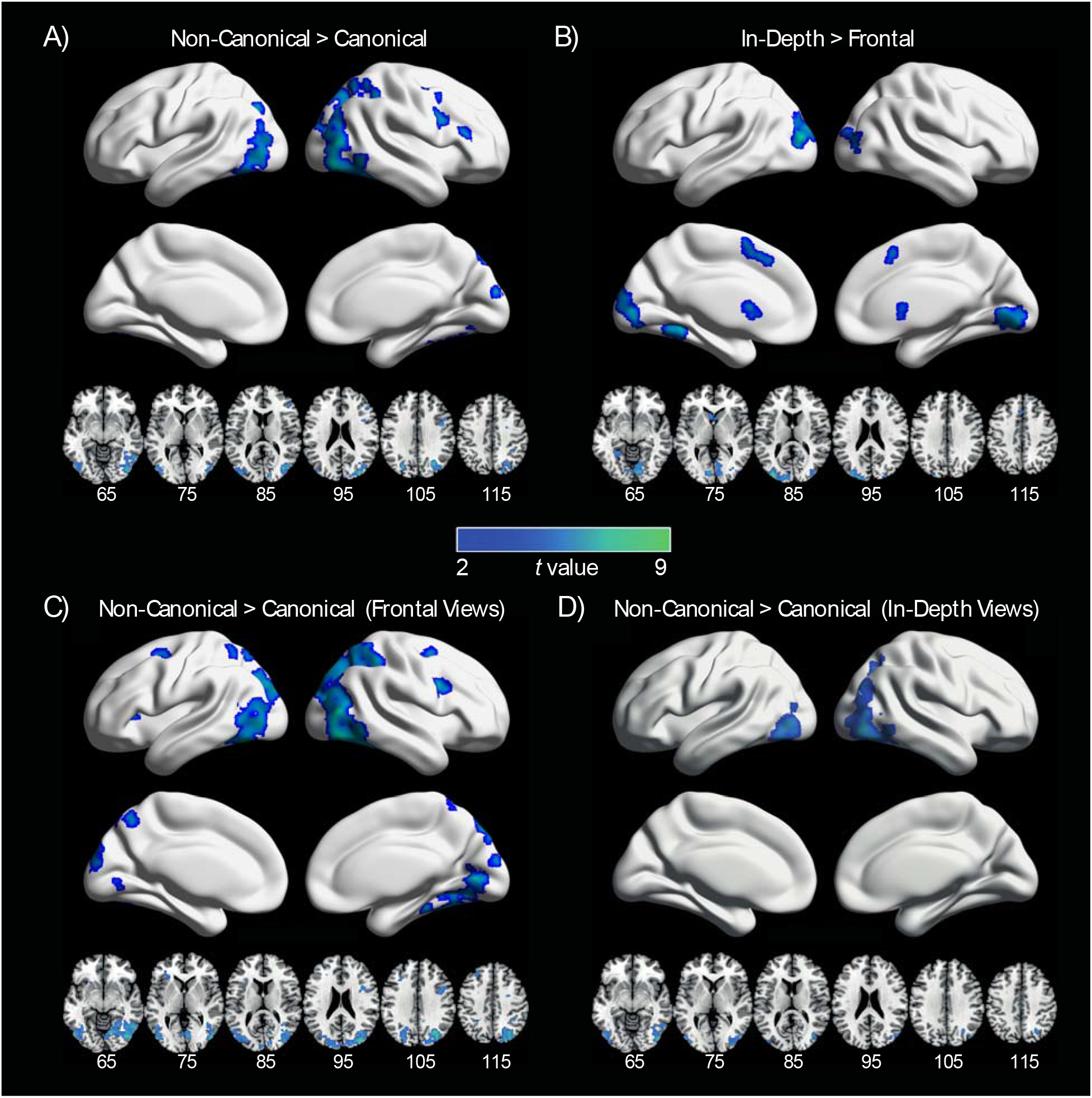
Results of the whole brain Analysis. The whole brain analysis was conducted in standard MNI space. Contrasts of interest were calculated within voxels significantly active throughout the whole experiment (Omnibus *F* contrast with *p* > 0.05 uncorr.) with a voxel-level threshold of *p* < 0.001 (uncorr.) and a cluster threshold of 100 voxels. Contrast images are displayed on a surface version of the ch2 brain using the BrainNet viewer (Xia, Wang, & He, 2013). **A)** Brain areas identified as showing higher activation for non-canonical compared to canonical objects (across in-depth and frontal views) were the lateral occipital complex (LOC) of both hemispheres, the fusiform gyri of both hemispheres as well as in the right supplementary motor cortex, the right parietal cortex, the right inferior frontal cortex and the right dorsolateral prefrontal cortex. **B)** Brain areas identified as showing higher activation for objects presented as in-depth rotated views compared to frontal views (across canonical and non-canonical views) were the occipital cortices of both hemispheres, the precentral cortices, frontal cortices of both hemispheres and the left fusiform gyrus. **C)** For objects presented in frontal views, higher activation for non-canonical compared to canonical objects were located in the parietal lobes, the LOC, premotor and supplementary motor cortices of both hemispheres, the right angular gyrus, the right fusiform, the right inferior frontal and the left frontal cortex. **D)** For objects presented as in-depth rotated views, higher activation for non-canonical compared to canonical objects were located in the LOC of both hemispheres and the right superior parietal lobe.

## Notes

### Competing Interest Statement

The authors have declared no competing interest.

## References

Alivisatos, B., & Petrides, M. (1997). Functional activation of the human brain during mental rotation. Neuropsychologia, 35(2), 111–118. https://doi.org/10.1016/S0028-3932(96)00083-8

Bálint, R. (1909). Seelenlähmung des “Schauens”, optische Ataxie, räumliche Störung der Aufmerksamkeit. Monatsschrift Für Psychiatrie Und Neurologie, 25(1), 51–66.

Balslev, D., Odoj, B., Rennig, J., & Karnath, H. (2014). Abnormal center-periphery gradient in spatial attention in simultanagnosia. Journal of Cognitive Neuroscience, 26(12), 2778–2788. https://doi.org/10.1162/jocn_a_00666

Best, F. (1917). Hemianopsie und Seelenblindheit bei Hirnverletzungen. Albrecht von Græfes Archiv für Ophthalmologie, 93(1), 49–150. https://doi.org/10.1007/BF01858120

Blanz, V., Tarr, M. J., & Bülthoff, H. H. (1999). What object attributes determine canonical views? Perception, 28(5), 575–599. https://doi.org/10.1068/p2897

Bloechle, J., Huber, S., Klein, E., Bahnmueller, J., Moeller, K., & Rennig, J. (2018). Neuro-cognitive mechanisms of global Gestalt perception in visual quantification. NeuroImage, /8/(April), 359–369. https://doi.org/10.1016/j.neuroimage.2018.07.026

Brainard, D. H. (1997). The Psychophysics Toolbox. Spatial Vision, 10(4), 433–436.

Bülthoff, H. H., Edelman, S. Y., & Tarr, M. J. (1995). How are three-dimensional objects represented in the brain? Cerebral Cortex (New York, N.Y.□: 1991), 5(3), 247–260.

Carlson, V. R. (1962). Size-constancy judgments and perceptual compromise. Journal of Experimental Psychology, 63(1), 68–73.

Chao, L. L., & Martin, A. (2000). Representation of Manipulable Man-Made Objects in the Dorsal Stream. NeuroImage, 12(4), 478–484. https://doi.org/10.1006/nimg.2000.0635

Cooper, A. C., & Humphreys, G. W. (2000). Coding space within but not between objects: Evidence from Balint’s syndrome. Neuropsychologia, 38(6), 723–733. https://doi.org/10.1016/S0028-3932(99)00150-5

Dale, A. M. (1999). Optimal experimental design for event-related fMRI. Human Brain Mapping, 8(2-3), 109–114.

Dale, A. M., Fischl, B., & Sereno, M. I. (1999). Cortical surface-based analysis. I. Segmentation and surface reconstruction. NeuroImage, 9(2), 179–194. https://doi.org/10.1006/nimg.1998.0395

Dekker, T., Mareschal, D., Sereno, M. I., & Johnson, M. H. (2011). Dorsal and ventral stream activation and object recognition performance in school-age children. Neuroimage, 57(3), 659–670. https://doi.org/10.1016/j.neuroimage.2010.11.005

Destrieux, C., Fischl, B., Dale, A., & Halgren, E. (2010). Automatic parcellation of human cortical gyri and sulci using standard anatomical nomenclature. NeuroImage, 53(1), 1–15. https://doi.org/10.1016/j.neuroimage.2010.06.010

Fischl, B., Salat, D. H., Busa, E., Albert, M., Dieterich, M., Haselgrove, C., van der Kouwe, A., Killiany, R., Kennedy, D., Klaveness, S., Montillo, A., Makris, N., Rosen, B., & Dale, A. M. (2002). Whole Brain Segmentation. Neuron, 33(3), 341–355. https://doi.org/10.1016/S0896-6273(02)00569-X

Freud, E., Culham, J. C., Plaut, D. C., & Behrmann, M. (2017). The large-scale organization of shape processing in the ventral and dorsal pathways. ELife, 6. https://doi.org/10.7554/eLife.27576

Freud, E., Ganel, T., Shelef, I., Hammer, M. D., Avidan, G., & Behrmann, M. (2017). Three-Dimensional Representations of Objects in Dorsal Cortex are Dissociable from Those in Ventral Cortex. Cerebral Cortex (New York, N.Y.: 1991), 27(1), 422–434. https://doi.org/10.1093/cercor/bhv229

Freud, E., Robinson, A. K., & Behrmann, M. (2018). More than Action: The Dorsal Pathway Contributes to the Perception of 3-D Structure. Journal of Cognitive Neuroscience, 30(7), 1047–1058. https://doi.org/10.1162/jocn_a_01262

Friedman-Hill, S. R., Robertson, L. C., & Treisman, A. (1995). Parietal contributions to visual feature binding: Evidence from a patient with bilateral lesions. Science (New York, N.Y.), 269(5225), 853–855. https://doi.org/10.1126/science.7638604

Goldhahn, D., Eckhart, T., & Quasthoff, U. (2012). Building Large Monolingual Dictionaries at the Leipzig Corpora Collection: From 100 to 200 Languages.

Goodale, M. A., Milner, A. D., Jakobson, L. S., & Carey, D. P. (1991). A neurological dissociation between perceiving objects and grasping them. Nature, 349(6305), 154–156. https://doi.org/10.1038/349154a0

Grill-Spector, K., Kushnir, T., Hendler, T., & Malach, R. (2000). The dynamics of object-selective activation correlate with recognition performance in humans. Nature Neuroscience, 3(8), 837–843. https://doi.org/10.1038/77754

Grill-Spector, Kalanit, Kourtzi, Z., & Kanwisher, N. (2001). The lateral occipital complex and its role in object recognition. Vision Research, 41(10-11), 1409–1422. https://doi.org/10.1016/S0042-6989(01)00073-6

Haxby, J. V., Gobbini, M. I., Furey, M. L., Ishai, A., Schouten, J. L., & Pietrini, P. (2001). Distributed and overlapping representations of faces and objects in ventral temporal cortex. Science (New York, N.Y.), 293(5539), 2425–2430. https://doi.org/10.1126/science.1063736

Haynes, J.-D., & Rees, G. (2006). Decoding mental states from brain activity in humans. Nature Reviews. Neuroscience, 7(7), 523–534. https://doi.org/10.1038/nrn1931

Himmelbach, M., Erb, M., Klockgether, T., Moskau, S., & Karnath, H.-O. (2009). FMRI of global visual perception in simultanagnosia. Neuropsychologia, 47(4), 1173–1177. https://doi.org/10.1016/j.neuropsychologia.2008.10.025

Huberle, E., & Karnath, H.-O. (2006). Global shape recognition is modulated by the spatial distance of local elements—Evidence from simultanagnosia. Neuropsychologia, 44(6), 905–911. https://doi.org/10.1016/j.neuropsychologia.2005.08.013

Huberle, E., & Karnath, H.-O. (2012a). The role of temporo-parietal junction (TPJ) in global Gestalt perception. Brain Structure & Function, 217(3), 735–746. https://doi.org/10.1007/s00429-011-0369-y

Huberle, E., & Karnath, H.-O. (2012b). The role of temporo-parietal junction (TPJ) in global Gestalt perception. Brain Structure and Function, 217(3), 735–746. https://doi.org/10.1007/s00429-011-0369-y

Jacobs, H. I. L., Gronenschild, E. H. B. M., Evers, E. A. T., Ramakers, I. H. G. B., Hofman, P. A. M., Backes, W. H., Jolles, J., Verhey, F. R. J., & Van Boxtel, M. P. J. (2015). Visuospatial processing in early Alzheimer’s disease: A multimodal neuroimaging study. Cortex; a Journal Devoted to the Study of the Nervous System and Behavior, 64, 394–406. https://doi.org/10.1016/j.cortex.2012.01.005

James, T. W., Humphrey, G. K., Gati, J. S., Menon, R. S., & Goodale, M. a. (2002). Differential effects of viewpoint on object-driven activation in dorsal and ventral streams. Neuron, 35(4), 793–801. https://doi.org/10.1016/S0896-6273(02)00803-6

Janssen, P., Verhoef, B.-E., & Premereur, E. (2018). Functional interactions between the macaque dorsal and ventral visual pathways during three-dimensional object vision. Cortex; a Journal Devoted to the Study of the Nervous System and Behavior, 98, 218–227. https://doi.org/10.1016/j.cortex.2017.01.021

Karnath, H. O., Ferber, S., & Bülthoff, H. H. (2000). Neuronal representation of object orientation. Neuropsychologia, 38(9), 1235–1241. https://doi.org/10.1016/S0028-3932(00)00043-9

Koffka, K. (1935). Principles of Gestalt Psychology (1st ed.). Harcourt.

Konen, C. S., & Kastner, S. (2008). Two hierarchically organized neural systems for object information in human visual cortex. Nature Neuroscience, 11(2), 224–231. https://doi.org/10.1038/nn2036

Kosslyn, S. M., Alpert, N. M., Thompson, W. L., Chabris, C. F., Rauch, S. L., & Anderson, a K. (1994). Identifying objects seen from different viewpoints. A PET investigation. Brain□: A Journal of Neurology, 117(5), 1055–1071. https://doi.org/10.1093/brain/117.5.1055

Kourtzi, Z., & Kanwisher, N. (2000). Cortical Regions Involved in Perceiving Object Shape. The Journal of Neuroscience, 20(9), 3310–3318. https://doi.org/10.1523/JNEUROSCI.20-09-03310.2000

Kristensen, S., Garcea, F. E., Mahon, B. Z., & Almeida, J. (2016). Temporal Frequency Tuning Reveals Interactions between the Dorsal and Ventral Visual Streams. Journal of Cognitive Neuroscience, 28(9), 1295–1302. https://doi.org/10.1162/jocn_a_00969

Landau, B., Hoffman, J. E., & Kurz, N. (2006). Object recognition with severe spatial deficits in Williams syndrome: Sparing and breakdown. Cognition, 100(3), 483–510. https://doi.org/10.1016/j.cognition.2005.06.005

Luria, A. (1959). Disorders of “simultaneous perception” in a case of bilateral occipito-parietal brain injury. Brain□: A Journal of Neurology, 82, 437–449.

Mahon, B. Z., Kumar, N., & Almeida, J. (2013). Spatial Frequency Tuning Reveals Interactions between the Dorsal and Ventral Visual Systems. Journal of Cognitive Neuroscience, 25(6), 862–871. https://doi.org/10.1162/jocn_a_00370

Martinaud, O., Mirlink, N., Bioux, S., Bliaux, E., Champmartin, C., Pouliquen, D., Cruypeninck, Y., Hannequin, D., & Gérardin, E. (2016). Mirrored and rotated stimuli are not the same: A neuropsychological and lesion mapping study. Cortex, 78, 100–114. https://doi.org/10.1016/j.cortex.2016.03.002

Moeller, S., Yacoub, E., Olman, C. A., Auerbach, E., Strupp, J., Harel, N., & Uğurbil, K. (2010). Multiband multislice GE-EPI at 7 tesla, with 16-fold acceleration using partial parallel imaging with application to high spatial and temporal whole-brain fMRI. Magnetic Resonance in Medicine, 63(5), 1144–1153. https://doi.org/10.1002/mrm.22361

Mruczek, R. E. B., von Loga, I. S., & Kastner, S. (2013). The representation of tool and non-tool object information in the human intraparietal sulcus. Journal of Neurophysiology, 109(12), 2883–2896. https://doi.org/10.1152/jn.00658.2012

Mumford, J. A., Turner, B. O., Ashby, F. G., & Poldrack, R. A. (2012). Deconvolving BOLD activation in event-related designs for multivoxel pattern classification analyses. NeuroImage, 59(3), 2636–2643. https://doi.org/10.1016/j.neuroimage.2011.08.076

Navon, D. (1977). Forest before trees: The precedence of global features in visual perception. Cognitive Psychology, 9(3), 353–383. https://doi.org/10.1016/0010-0285(77)90012-3

Pelli, D. G. (1997). The VideoToolbox software for visual psychophysics: Transforming numbers into movies. Spatial Vision, 10(4), 437–442.

Rennig, J., Bilalić, M., Huberle, E., Karnath, H.-O., & Himmelbach, M. (2013a). The temporo-parietal junction contributes to global gestalt perception-evidence from studies in chess experts. Frontiers in Human Neuroscience, 7, 513. https://doi.org/10.3389/fnhum.2013.00513

Rennig, J., Bilalić, M., Huberle, E., Karnath, H.-O., & Himmelbach, M. (2013b). The temporo-parietal junction contributes to global gestalt perception—Evidence from studies in chess experts. Frontiers in Human Neuroscience, 7. https://doi.org/10.3389/fnhum.2013.00513

Rennig, J., Himmelbach, M., Huberle, E., & Karnath, H.-O. (2015). Involvement of the TPJ area in processing of novel global forms. Journal of Cognitive Neuroscience, 27(8), 1587–1600. https://doi.org/10.1162/jocn_a_00809

Rennig, J., & Karnath, H.-O. (2016). Stimulus size mediates Gestalt processes in object perception— Evidence from simultanagnosia. Neuropsychologia, 89, 66–73. https://doi.org/10.1016/j.neuropsychologia.2016.06.002

Schendan, H. E., & Stern, C. E. (2007). Mental rotation and object categorization share a common network of prefrontal and dorsal and ventral regions of posterior cortex. NeuroImage, 35(3), 1264–1277. https://doi.org/10.1016/j.neuroimage.2007.01.012

Schendan, H. E., & Stern, C. E. (2008). Where vision meets memory: Prefrontal-posterior networks for visual object constancy during categorization and recognition. Cerebral Cortex (New York, N.Y.: 1991), 18(7), 1695–1711. https://doi.org/10.1093/cercor/bhm197

Shepard, R. N., & Metzler, J. (1971). Mental rotation of three-dimensional objects. Science (New York, N.Y.), 171(3972), 701–703. https://doi.org/10.1126/science.171.3972.701

Solms, M., Kaplan-Solms, K., Saling, M., & Miller, P. (1988). Inverted vision after frontal lobe disease. Cortex; a Journal Devoted to the Study of the Nervous System and Behavior, 24(4), 499–509.

Sugio, T., Inui, T., Matsuo, K., Matsuzawa, M., Glover, G. H., & Nakai, T. (1999). The role of the posterior parietal cortex in human object recognition: A functional magnetic resonance imaging study. Neuroscience Letters, 276(1), 45–48.

Terhune, K. P., Liu, G. T., Modestino, E. J., Miki, A., Sheth, K. N., Liu, C.-S. J., Bonhomme, G. R., & Haselgrove, J. C. (2005). Recognition of objects in non-canonical views: A functional MRI study. Journal of Neuro-Ophthalmology: The Official Journal of the North American Neuro-Ophthalmology Society, 25(4), 273–279.

Turnbull, O. H., Beschin, N., & Della Sala, S. (1997). Agnosia for object orientation: Implications for theories of object recognition. Neuropsychologia, 35(2), 153–163.

Turnbull, O. H., Laws, K. R., & McCarthy, R. A. (1995). Object recognition without knowledge of object orientation. Cortex; a Journal Devoted to the Study of the Nervous System and Behavior, 31(2), 387–395.

Ungerleider, L., & Mishkin, M. (1982). Two cortical visual systems. In D. J. Ingle, M. A. Goodale, & R. J. E. Mansfield (Eds.), Analysis of visual behavior (pp. 549–586). MIT Press.

Van Dromme, I. C., Premereur, E., Verhoef, B.-E., Vanduffel, W., & Janssen, P. (2016). Posterior Parietal Cortex Drives Inferotemporal Activations During Three-Dimensional Object Vision. PLOS Biology, 14(4), e1002445. https://doi.org/10.1371/journal.pbio.1002445

Verfaillie, K., & Boutsen, L. (1995). A corpus of 714 full-color images of depth-rotated objects. Perception & Psychophysics, 57(7), 925–961.

Wagemans, J., Elder, J. H., Kubovy, M., Palmer, S. E., Peterson, M. a., Singh, M., & von der Heydt, R. (2012). A century of Gestalt psychology in visual perception: I. Perceptual grouping and figureground organization. Psychological Bulletin, 138(6), 1172–1217. https://doi.org/10.1037/a0029333

Wagemans, J., Feldman, J., Gepshtein, S., Kimchi, R., Pomerantz, J. R., van der Helm, P. A., & van Leeuwen, C. (2012). A century of Gestalt psychology in visual perception: II. Conceptual and theoretical foundations. Psychological Bulletin, 138(6), 1218–1252. https://doi.org/10.1037/a0029334

Weissman, D. H., & Woldorff, M. G. (2005). Hemispheric asymmetries for different components of global/local attention occur in distinct temporo-parietal loci. Cerebral Cortex (New York, N.Y.□: 1991), 15(6), 870–876. https://doi.org/10.1093/cercor/bhh187

Wertheimer, M. (1923). Untersuchungen zur Lehre von der Gestalt. II. Psychologische Forschung, 4(1), 301–350. https://doi.org/10.1007/BF00410640

Wolpert, I. (1924). Die Simultanagnosie—Störung der Gesamtauffassung. Zeitschrift Für Die Gesamte Neurologie Und Psychiatrie, 93, 397–415.

Xia, M., Wang, J., & He, Y. (2013). BrainNet Viewer: A network visualization tool for human brain connectomics. PloS One, 8(7), e68910. https://doi.org/10.1371/journal.pone.0068910

Zaretskaya, N., Anstis, S., & Bartels, A. (2013). Parietal cortex mediates conscious perception of illusory gestalt. The Journal of Neuroscience □: The Official Journal of the Society for Neuroscience, 33(2), 523–531. https://doi.org/10.1523/JNEUROSCI.2905-12.2013

